# Association cortical areas in the mouse contain a large population of fast-spiking GABAergic neurons that do not express parvalbumin

**DOI:** 10.1101/2023.11.02.565290

**Authors:** Erik Justin Courcelles, Kasper Kjelsberg, Laura Convertino, Rajeevkumar Raveendran Nair, Menno P. Witter, Maximiliano José Nigro

## Abstract

GABAergic neurons represent 10-15% of the neuronal population of the cortex but exert a powerful control over information flow in cortical circuits. The largest GABAergic class in the neocortex is represented by the parvalbumin-expressing (PV-INs) fast-spiking neurons, which provide powerful somatic inhibition to their postsynaptic targets. Recently, the density of PV-INs has been shown to be lower in associative areas of the mouse cortex as compared to sensory and motor areas. Modeling work based on these quantifications linked the low-density of PV-INs with specific computations of associative cortices. However, it is still unknown whether the total GABAergic population of association cortices is smaller or whether another GABAergic type can compensate for the low density of PV-INs. In the present study we investigated these hypotheses using a combination of neuroanatomy, mouse genetics and neurophysiology. We found that the GABAergic population of association areas is comparable to that of primary sensory areas, and it is enriched of fast-spiking neurons that do not express PV and were not accounted for by previous quantifications. We developed an intersectional viral strategy to demonstrate that the synaptic output of fast-spiking neurons is comparable across cortical regions. Our results provide quantifications of the density and output strength of fast-spiking GABAergic neurons and offers new biological constrains to refine current models of cortical computations.

## Introduction

The wide neuronal diversity of the cerebral cortex is thought to endow it with a variety of circuit motifs that shape information processing (Harris and Shepherd 2015; Luo, 2021). The cerebral cortex can be parcellated into functional areas that show specific cytoarchitectonic features and expression of molecular markers (van Essen and Glasser, 2018). The relationship between cortical parcellation and distribution of neuron types in the cortex is poorly understood. Recent studies unveiled a correlation between the distribution of GABAergic types and the connectivity-based hierarchical organization of the cortex (Kim et al., 2017; Ding et al., 2023). GABAergic neurons represent 10-15% of the neuronal population and their diversity allows them to participate in different cortical microcircuits (Kepecs and Fishell, 2014; Tremblay et al., 2016). GABAergic neurons are divided in two major groups according to embryonal origin: those derived from the medial ganglionic eminence (MGE) and those derived from the caudal ganglionic eminence (CGE) (Rudy et al., 2011). MGE-derived GABAergic neurons are further divided according to molecular marker expression into parvalbumin (PV)-expressing and somatostatin (SST)-expressing interneurons (INs) (Tremblay et al., 2016). CGE-derived interneurons (CGE-INs) have been divided in VIP-expressing and VIP non-expressing neurons (Lee et al., 2010). Transcriptomic analysis has confirmed this classification and further divided the CGE group into three major classes: vasoactive intestinal peptide (VIP)-expressing, Lamp5, and Sncg groups (Tasic et al., 2018).

Molecularly defined GABAergic neurons have been shown to be homogeneously distributed across sensory-motor areas of the neocortex (Xu et al., 2010). However, recent cortex-wide examination of the distribution of three major GABAergic groups (PV, SST, VIP) revealed a reduction of the density of PV-INs in association cortices (Whissell et al., 2015; Kim et al., 2017). PV-INs are the most abundant GABAergic type in the neocortex, they show a characteristic fast-spiking firing pattern and provide powerful perisomatic inhibition to their postsynaptic targets (Tremblay et al., 2016). In sensory cortices, PV-INs participate in feedforward inhibitory circuits gating information flow in the thalamocortical circuit and in translaminar circuits (Pouille and Scanziani, 2001; Gabernet et al., 2005; Helmstaedter et al., 2008). In association cortices, PV-INs have been shown to receive long-range excitatory inputs and have been shown to mediate feedforward inhibition (McGarry and Carter, 2016; Anastasiades et al., 2018; Willems et al., 2018). Their role in sensory gating through feedforward inhibition led to the hypothesis that PV-INs control the flow of sensory information in the cortico-hippocampal network (DeCurtis and Paré, 2004; Willems et al., 2018). This is in striking contrast with the low density of PV-INs in association areas described previously (Whissell et al., 2015; Kim et al., 2017). The low density of PV-INs suggests that either the GABAergic population of association areas is smaller than that of the sensory-motor areas, or that other GABAergic cell-types are enriched. In the present study we aimed to explore these two scenarios by combining mouse genetics, molecular marker expression, enhancer mediated transgene expression and electrophysiological characterization of GABAergic neurons in the cortex. We found that indeed the GABAergic population of association areas is not smaller than in sensory cortices, suggesting an enrichment of other interneuron types. We identified a large group of fast-spiking neurons that do not express PV in association cortex and developed an intersectional viral approach to label them, which allowed us to show that the density of fast-spiking neurons is comparable across cortical regions. Using this approach, we were able to measure their synaptic output onto excitatory neurons and demonstrate that it is also comparable to that of fast-spiking neurons in sensory cortices.

Our results unveiled the presence of a large GABAergic population that was not accounted for because of the lack of markers and tools for labeling it. Our study provides new constrains to guide future modeling work about the computational abilities of different cortical regions.

## Materials and methods

### Animal models

The research described here was performed on adult mice (>8 weeks old) of both sexes. We used the following mouse lines from Jackson Laboratories: C57BL/6J mice (n= 6) (stock 000664), SST-Cre mice (n= 6) (stock 013044), PV-cre mice (stock 008069), and R26R-ChR2-YFP (n= 9) (stock 012569). The GAD67-GFP mice (n= 23) were obtained as gift from Dr. Yanagawa (Tamamaki et al., 2003). The GAD67-GFP and SST-cre mice were bred as heterozygous. The PV-cre mice were crossed with the R26R-ChR2-YFP mice (stock 012569) to obtain the PV-ChR2 mice (n= 9). Animals were bred in house, group housed in enriched cages with water and food ad libitum. Animals were kept with an inverted dark/light cycle of 12h/12h.

### Adeno-associated virus (AAV) production

For producing AAV1-S5E2-dTom (S5E2-dTom), the plasmid construct pAAV-S5E2-dTom was purchased from Addgene (#135630) (Vormstein-Schneider et al., 2020). For AAV1-S5E2-lox2272 dTom loxP-reverse complement (rc)GFP lox2272 loxp (S5E2-*lox* dTom *lox* rcGFP *lox2*) and AAV1-S5E2-loxP ChR2 (H134R)-mCherry loxP (S5E2-*lox* ChR2 mCh *lox*), first we synthesized the corresponding gene segments and cloned in the pAAV-S5E2-dTom plasmid backbone as described below. Briefly, using NheI and EcoRV restriction sites in pAAV-S5E2-dTom plasmid we cloned in the sequence of nls-dTomato flanked by incompatible lox2272 and loxP sites followed by reverse complement sequence of GFP followed by two lox sites lox2272 and loxP sites at the 3’ end. This arrangement of lox sites and gene sequences driven by S5E2 promoter in S5E2-*lox* dTom *lox* rcGFP *lox2* enable the expression of dTomato in E2/SST Cre-cells while the expression of GFP only after Cre-mediated recombination (in E2/SST Cre+ cells). For the S5E2-*lox* ChR2 mCh *lox* plasmid production, loxP-ChR2 (H134R)-mCherry-loxP gene segment was cloned into the pAAV-S5E2-dTom using NheI and EcoRV restriction sites. The expression of ChR2-mCherry will be driven by S5E2 promoter, but it will not express in Cre+ cells.

S5E2-*lox* dTom *lox* rcGFP *lox2* and S5E2-*lox* ChR2 mCh *lox* were packaged in AAV1 and purified using iodixanol density gradient method. Specifically, the corresponding pAAV plasmid construct along with AAV helper plasmids encoding the structural elements were transfected using polyethylenimine (PEI) into the AAV-293 cell line (CVCL_6871, Agilent, USA). The day before transfection, 7 x 10^6^ AAV-293 cells were seeded into 150 mm cell culture plates in DMEM (# 41965062, Thermo Fisher Scientific) containing 10% fetal bovine serum (#16000-044, ThermoFisher, USA) and penicillin/streptomycin antibiotics (#15140122, Thermo Fisher Scientific). All the plasmids for AAV vector preparations were made using endotoxin free plasmid maxiprep kit (#12663, Qiagen). PEI mediated co-transfection of pAAV plasmid-containing the transgene, pHelper and pAAV1 capsid helper plasmids were carried out on next day. After 24 hours, the medium was replaced with fresh 10% FBS-containing DMEM. The AAV-293 cells were cultured for two days following transfection to allow AAV synthesis to occur in cells. The medium and AAV-293 cells filled with virus particles were scraped from the cell culture plates, then isolated by centrifugation at 200 x g. The cell pellet was then subjected to lysis using 20 mM Tris, 300 mM NaCl and 20 mM MgCl_2_, pH 7.6 buffer. The supernatant was mixed with 40% Polyethylene Glycol 8000 (PEG) for 2 hours in ice for precipitation of virus particles. The PEG treated medium was centrifuged at 4000 x g for 15 minutes. The lysate and the PEG-precipitate was treated with benzonase nuclease HC (#71206-3, Millipore) for 45 minutes at 37°C. Benzonase-treated lysate was centrifuged at 3000 x g for 15 mins and the clear supernatant then subjected to iodixanol gradient ultracentrifugation. 4 different layers of the gradient, i.e., 15%, 25%, 40%, and 58% of iodixanol was built in a Beckman quick-seal centrifuge tube. Phenol red was added to the 25% and 58% layers to aid visualization of the layers within the tube. The virus containing supernatant was layered above the 15% iodixanol by slowly dripping the solution onto the top layer of the gradient. Seal the tip of the tube using a heating device (Beckman and Coulter). Sealed tubes were centrifuged at 350,000 g for 90 min in a T70i rotor at 10°C. The 40% iodixanol layer after ultracentrifugation was collected and buffer exchanged with DPBS using Amicon Ultra centrifugal filters (#Z648043, Millipore) (modified protocol from Addgene, USA). We performed quantitative PCR on the viral stocks and the titer was determined as approximately 1.9 x 10^12^ genomic copies/ml for S5E2-*lox* dTom *lox* rcGFP *lox2* and 1.1 x 10^12^ genomic copies/ml for S5E2-*lox* ChR2 mCh *lox*.

### Injections of viral vectors

The AAVs were injected into the brains of C57 (n= 6), and SST-cre (n= 3) mice. Mice were anaesthetized with isoflurane (4% Nycomed, airflow 1 l/min) in an induction chamber. The mice were placed in a stereotactic frame on a heated pad (37°C) throughout the procedure and head-fixed with ear bars. Eye ointment was applied to protect the cornea. The following analgesics were applied subcutaneously: buprenorphine hydrochloride (0.1 mg/Kg, Temgesic, Invidior), meloxicam (1 mg/Kg, Metacam Boerringer Ingelheim Vetmedica), bupivacaine hydrochloride (1 mg/Kg locally, Marcain, Astra Zeneca).

The skin on the incision site was shaved and disinfected with Pyrisept. An incision was made to access the skull. The skull was thinned with a dental drill at the desired location, and a hole was punched with the tip of a glass pipette. We used the following coordinates from bregma: for wS1 −0.7 mm AP, +3.1 mm ML, −0.15 and −0.55 DV; for PER/ECT −3 mm AP, −1.5 DV, for the ML the pipette was moved to the edge of the skull and then retracted medially 0.4 mm; for PrL/IL: +1.8 mm AP, 0.25 mm ML, −1.3 mm DV. Injections of 100-200 nl (50 nl/minute) in each location were made with a glass pipette (20-30 µm tip size) attached to an injector (Nanoliter 2010, World Precision Instruments) controlled by a microsyringe pump controller (Micro4 pump, World Precision Instruments). After retracting the pipette, the wound was rinsed with saline and sutured. The mice were allowed to recover in a heated chamber (37°C) before returning to the homecage. Postoperative analgesic was applied 6-8 h after the procedure (Temgesic) and 24 h after the procedure (Metacam). The survival time for transduction and expression of the genetic material was 10-15 days.

### In vitro electrophysiology

Mice of either sex (>8 week old) were euthanized with an overdose of pentobarbital (i.p. 100 mg/Kg, Apotekerforeninger), before intracardial perfusion with cutting solution (RT) of the following composition (in mM): 93 choline chloride, 3 KCl, 1.25 NaH_2_PO_4_, 30 NaHCO_3_, 20 HEPES, 10 glucose, 5 MgCl_2_, 0.5 CaCl_2_, 5 N-acetylcysteine, saturated with 95% O_2_ and 5% CO_2_. The brain was extracted and a section containing either PER or wS1 was glued onto the stage of a vibratome (VT1000, Leica) filled with cold (4°C) cutting solution. Slices (300 µm) were transferred to a chamber filled with warm (34°C) cutting solution and incubated for 15 minutes. The slices were stored in a different chamber filled with the following solution (in mM) (RT): 92 NaCl, 3 KCl, 1.25 NaH_2_PO_4_, 30 NaHCO_3_, 20 HEPES, 10 glucose, 5 MgCl_2_, 0.5 CaCl_2_, 5 N-acetylcysteine, saturated with 95% O_2_ and 5% CO_2_.

The slices were placed in a recording chamber of an upright microscope (BW51, Olympus) filled with warm (35°C) recording solution of the following composition: 124 NaCl, 3 KCl, 1.25 NaH_2_PO_4_, 26 NaHCO_3_, 10 glucose, 1 MgCl_2_, 1.6 CaCl_2_, saturated with 95% O_2_ and 5% CO_2_. Characterization of the electrophysiological properties was performed in presence of synaptic blockers: 10 µM DNQXNa2, 25 µM APV and 10 µM gabazine. wS1 was identified under DIC optics by the presence of characteristic barrels in layer 4. To identify A35 under DIC optics, we first identified the lateral entorhinal cortex (LEC) by its characteristic large neurons in layer 2a, and by the clear separation between layer 2a and 2b. A35 was defined as the cortex dorsally located to LEC. Only healthy, fluorophore-expressing neurons were selected for recordings and no other criterion was applied. Whole-cell patch clamp recordings were performed with borosilicate pipettes (Sutter Instruments) with a resistance of 3-7 MΩ filled with a solution of the following composition (in mM): 130 K-gluconate, 10 KCl, 10 HEPES, 0.2 EGTA, 4 ATP-Mg, 0.3 GTP-Na, 5 phosphocreatine-Na2, pH 7.3. In all recordings biocytin was added to the pipette solution to recover the layer location of the neuron. Membrane potentials reported were not corrected for a junction potential of −14 mV. Current clamp recordings were performed with a Multiclamp 700B amplifier (Molecular Devices), digitized with a 1550A Digidata (Molecular Devices), interfaced with a personal computer with pClamp 11 (Molecular Devices). Data were sampled at 40 kHz and low-passed filtered at 10 kHz. All recordings were performed from a holding potential of −65 mV. The electrophysiological parameters analyzed were defined as follows: Input resistance (IR, MΩ): resistance measured from Ohm’s law from the peak of voltage responses to −25 pA hyperpolarizing current steps.

Sag ratio (dimensionless): measured from voltage responses peaking at −90 ± 2 mV to hyperpolarizing current steps. It is defined as the ratio between the voltage at the steady-state response and the peak. Rheobase (Rheo, pA): current level that evoked the first spike during a 1-s-long depolarizing ramp (500 pA/s).

Action potential threshold (APthr, mV): measured from action potentials (AP) evoked near rheobase, and defined as the voltage where the rise of the AP was 20 mV/ms.

AP half width (AP_HW_, ms): duration of the AP at half amplitude from APthr, measured from action potentials (AP) evoked near rheobase.

Amplitude of the fast afterhyperpolarization (fAHP, mV): measured from APthr, from action potentials (AP) evoked near rheobase.

AHP duration (AHPdur, ms): measured from action potentials (AP) evoked near rheobase and defined as the time difference between the fAHP and the most depolarized voltage in a 200 ms window. Maximal firing frequency (Fmax, AP/s): maximal firing frequency evoked with 1-s-long depolarizing current steps.

Voltage clamp experiments were performed from 9 PV-cre/ChR2 mice and 1 SST-cre mouse injected with the intersectional virus expressing channelrhodopsin (ChR2). The neurons were recorded using a pipette solution of the following composition (in mM): 130 CsMeSO_3_, 5 CsCl, 10 HEPES, 0.2 EGTA, 4 Mg-ATP, 0.3 Na-GTP, 5 Phosphocreatine-Na_2_, 5 QX-314-Cl. The extracellular solution (same as above) contained 10 µM DNQXNa2, 25 µM APV to block fast glutamatergic transmission. Inhibitory post-synaptic currents (IPSCs) were recorded at a holding potential of +10 mV. IPSCs were evoked by full field 470 nm light stimulation of the duration of 1 ms, delivered through a 40X objective. Recordings were not corrected for a 10 mV junction potential.

### Histology

Mice used for histological analysis were anaesthetized with isoflurane and then euthanized with an overdose of pentobarbital (i.p. 100 mg/Kg, Apotekerforeninger). The mice were intracardially perfused with Ringer solution (0.85% NaCl, 0.025% KCl, 0.02% NaHCO3) followed by 4% paraformaldehyde (PFA) in phosphate buffer (PB) (pH 7.4). The brain was extracted from the skull and placed in 4% PFA for 3 hours before moving it to a solution containing 15% sucrose in PB overnight. The brains were then stored in 30% sucrose for two days before being sliced at 50 µm with a freezing microtome. Slices were collected in six equally spaced series and placed in tubes containing a solution composed of 30% glycerol, 30% ethylene glycol and 40% phosphate buffer saline (PBS). The tissue was stored at −20 °C until used for histology.

The slices were washed in PB (3 × 10 minutes) before incubation with blocking solution (0.1% Triton-X, 10% NGS in PB). After blocking the tissue was incubated with the primary antibody for three days at 4°C in a solution containing: 0.1% Triton-X, 1% NGS in PB. After being washed (3 × 1 hour in PB), the tissue was incubated with the secondary antibody overnight at 4 °C. The slices were washed (3 × 10 minutes in PB) and mounted on SuperFrost slides (Termo Fisher Scientific) and left to dry overnight. The slides were coversliped with entellan xylene (Merck Chemicals, Darmstadt, Germany) after being washed in xylene for 15 minutes, or with Fluoromount after being washed in distilled water for about 30 seconds. We used the following primary antibodies: Guinea Pig IgG anti-NeuN (1:1000, Sigma Millipore, #ABN90P), Rabbit anti-parvalbumin (1:1000, Swant, #PV-27), Mouse IgG1 anti-parvalbumin (1:1000, Sigma, #P3088), Rat IgG2a anti-RFP (1:1000, Proteintech, #5f8), Chicken IgY anti-GFP (1:1000, Abcam, #ab13970), Rabbit IgG anti-SST (1:1000, BMA Biomedicals),. We used the following secondary antibodies: Goat anti-guinea pig (IgG H+L) A647 (1:500, Invitrogen, #A-21450), Goat anti-Rat (IgG H+L) A-546 (1:500, Invitrogen, #A11081), Goat anti-Rabbit (IgG H+L) A488 (1:500, Invitrogen, #A11008), Goat anti-chicken (IgY H+L) A488 (1:500, Invitrogen, #A11039), Goat anti-rabbit (IgG H+L) A546 (1:500, Invitrogen, #A11010), Goat anti-rabbit (IgG H+L) A635 (1:500, Invitrogen, A31576).

### Image acquisition and analysis

We used a confocal microscope (Zeiss LSM 880 AxioImager Z2) to image regions of interest (ROIs) to count labeled neurons. We imaged ROIs including the following cortical areas: primary somatosensory cortex whisker field (wS1), primary visual cortex (V1), perirhinal cortex (PER), ectorhinal cortex (ECT), prelimbic cortex (PrL) and infralimbic cortex (IL). Images of ROIs were taken with a x20 air objective with 1 Airy unit pinhole size and saved as czi files. We selected 2 to 5 ROIs for each cortical region. ROIs contained the whole cortical area of interest in a given slice. Files containing the ROIs were uploaded in Neurolucida (Micro Bright Field Bioscience) for analysis. We created contours to delineate the layers of the ROIs using the PV signal: layer 4 in sensory areas was identified as the layer with the densest staining bordered by layer 2/3 superficially and layer 5A as a layer with low density of PV staining. PER and ECT were delineated according to Beaudin et al. (2013). We use here the nomenclature of PER (perirhinal) and ECT (ectorhinal) as reported in the Allen Brain Atlas and in the Franklin and Paxinos (2007) atlas. These two cortical regions correspond to A35 and A36 of the nomenclature used in Beaudin et al. (2013) and in several other reports on different rodent and primate species. We opted to follow the nomenclature of the Allen Brain Atlas to allow for easier comparison with previous quantifications of inhibitory cell types in the cortex (Whissell et al., 2015; Kim et al., 2017). PrL and IL were delineated according to Van De Werd et al. (2010). These delineations agreed with cytoarchitectural features obtained from the NeuN signal from adjacent slices. Symbols for each signal were used to count cells labeled by different markers. We counted all cells contained within the contours. Since we counted slices distanced 300 µm from each other, overcounting in the z axis is not relevant and we did not apply any correction. Quantification of the number of markers in each contour was done in Neurolucida Explorer (Micro Bright Field Bioscience) and exported in Microsoft Excel. The density of neurons labeled by a marker was measured as the number of labeled neurons in a contour divided by the area of the contour (cells/mm2). The GAD67-GFP mouse line labeled virtually all PV-INs but showed a lower specificity for SST-INs across all cortical areas included in our study (Fig. 1A-P; Table 1). For this reason, the number of PV-INs, and SST-INs was measured from the immunolabelling for the specific marker, independently of GFP expression. Transcriptomic analysis of the neuronal population of the cortex shows that SST, and PV are expressed exclusively in GABAergic neurons. We excluded from the counting a small population of PV-IR pyramidal neurons in L5B of wS1 (van Brederode et al., 1991). Values are reported as mean ± standard deviation). All graphs were created in Microsoft Excel or Matlab 2021a (Mathworks), images of ROIs were created in Zen lite (Zeiss) and figures in Adobe Illustrator.

**Figure 1.**
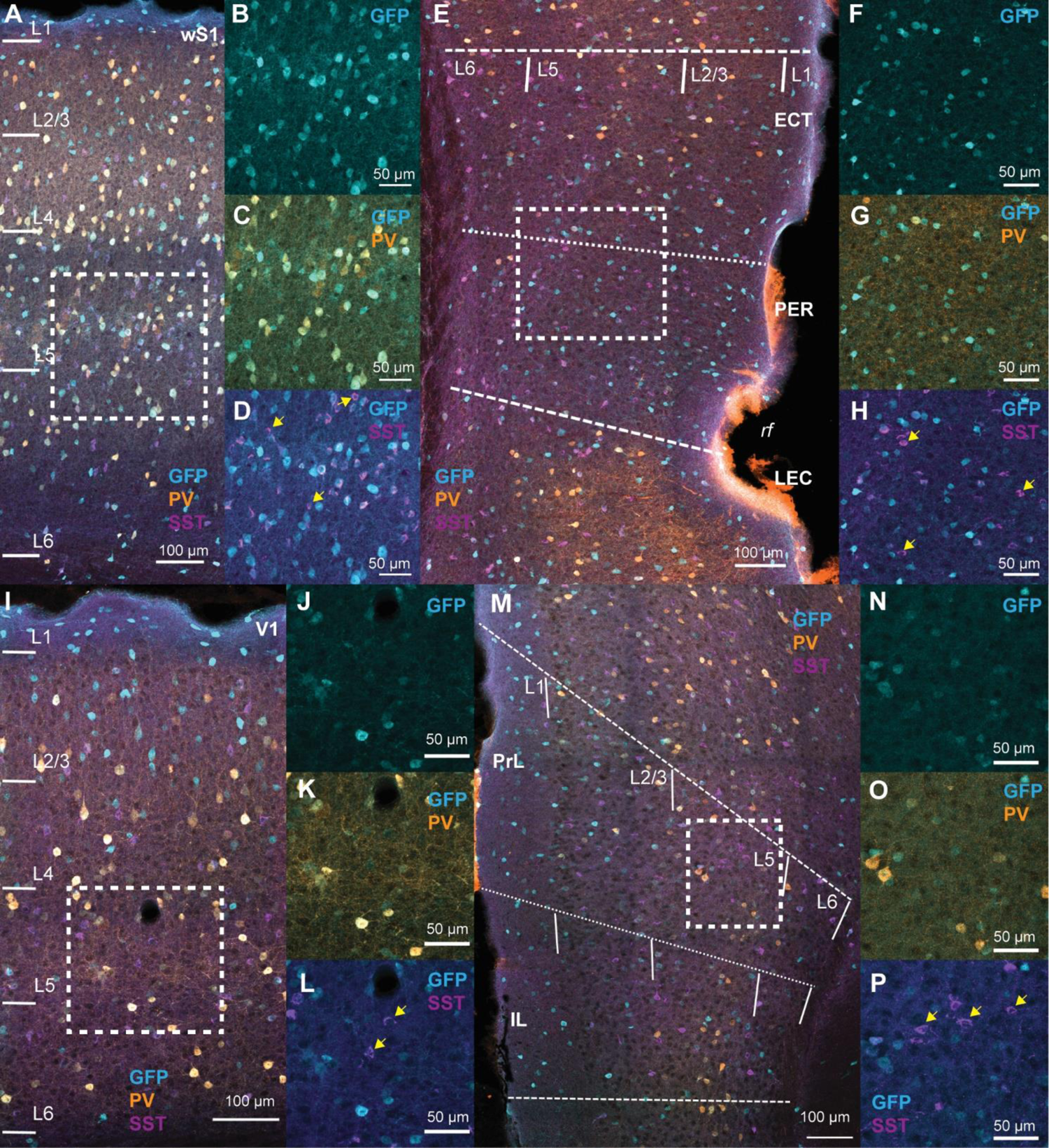
Efficiency of the GAD67-GFP mouse line to label PV and SST neurons. **A.** Representative confocal stack of a section of wS1 showing the overlap of the signals for GFP (turquoise), PV (orange) and SST (violet). **B-D.** Magnified pictures of the area enclosed in the white square in A showing the signals for GFP (B), GFP and PV (C) and GFP and SST (D). Arrows in D point at cells expressing SST but not GFP. **E-H.** Same as in A-D but for PER/ECT. H shows that also in PER/ECT some SST neurons are not labeled by GFP (arrows). LEC, lateral entorhinal cortex; rf, rhinal fissure. **I-L.** Same as for A-D but for V1. L shows that also in V1 some SST neurons do not express GFP (arrows). **M-P.** Same as for A-D but for PrL/IL. P shows that in PrL/IL some SST neurons do not express GFP.

**Table 1.** Efficiency of the GAD67 mouse line to label SST-INs and PV-INs in the six cortical areas included in this study. Rows represent the layers in each cortical area. Values in columns represent the mean and standard deviation (in parentheses) of the number of SST-INs, PV-INs and the efficiency for each. The mouse line consistently showed a slightly lower efficiency for SST-INs.

| Layer /<br>Cortex | SST (cells)<br>(n= 3 mice) | SST/GFP+<br>(cells)<br>(n= 3 mice) | Efficiency<br>(%)<br>(n= 3 mice) | PV (cells)<br>(n= 3 mice) | PV/GFP+ (cells)<br>(n= 3 mice) | Efficiency (%)<br>(n= 3 mice) |
| --- | --- | --- | --- | --- | --- | --- |
| L1 / V1 | 3 (0) | 2 (1) | 66.7 (33.3) | 1.7 (2.1) | 0.3 (0.6) | 12.5 (17.7) |
| L2/3 / V1 | 58 (2.7) | 52 (2) | 89.9 (6.9) | 118.7 (7.2) | 117.3 (7.6) | 98.9 (0.5) |
| L4 / V1 | 78.7 (18.5) | 66 (18.1) | 83.6 (9.9) | 128.7 (31.8) | 128.3 (32.3) | 99.6 (0.6) |
| L5 / V1 | 142.7 (37.2) | 118 (36.5) | 82.5 (9.6) | 194 (44.9) | 193 (43.3) | 99.6 (0.7) |
| L6 / V1 | 94.67 (24.03) | 74 (22.9) | 77.9 (9) | 115.7 (31.4) | 115.6 (31.4) | 100 (0) |
| L1 / wS1 | 2 (2.6) | 2 (2.7) | 100 (0) | 0 (0) | 0 (0) |  |
| L2/3 / wS1 | 56 (7.2) | 42.3 (1.5) | 76.2 (7.6) | 113.3 (10.5) | 112 (10) | 98.8 (0.4) |
| L4 / wS1 | 68 (4.4) | 45.3 (7.5) | 67.3 (15.5) | 221 (42.6) | 220.7 (42.0) | 99.9 (0.2) |
| L5 / wS1 | 142.7 (32) | 119.7 (25.4) | 84.0 (3.6) | 234.3 (42.2) | 234.3 (42.2) | 100 (0) |
| L6 / wS1 | 118.3 (19.6) | 85 (6.2) | 72.6 (6.9) | 144.7 (14.6) | 144.3 (14.0) | 99.8 (0.4) |
| L1 / A36 | 2.7 (2.3) | 2.7 (2.3) | 100 (0) | 0.7 (0.6) | 0.3 (0.6) | 100 (0) |
| L2/3 / A36 | 72.3 (5.8) | 60.3 (11.7) | 83.8 (8.7) | 46 (19.9) | 45.3 (20.5) | 99.6 (2.4) |
| L5 / A36 | 137 (26.1) | 106 (37.0) | 78.8 (8.8) | 103.7 (34.5) | 102.7 (35.1) | 99.0 (2) |
| L6 / A36 | 56.3 (7.1) | 49.7 (8.1) | 89.3 (4.7) | 43.7 (6.0) | 43.3 (6.5) | 100 (0) |
| L1 / A35 | 2.7 (0.6) | 2.7 (0.6) | 100 (0) | 0.3 (0.7) | 0.3 (0.6) | 100 (0) |
| L2/3 / A35 | 74.7 (14.6) | 60 (16.1) | 80.9 (7.9) | 23.3 (13.7) | 22.7 (14.2) | 99.8 (0.4) |
| L5 / A35 | 93.7 (15.9) | 73.3 (12.0) | 78.3 (5.6) | 55.3 (32.9) | 55.3 (32.9) | 98.8 (2.4) |
| L6 / A35 | 51.7 (5.5) | 44.7 (3.8) | 89.6 (4.3) | 26 (14) | 26 (14) | 100 (0) |
| L1 / PrL | 7.7 (4.0) | 5.3 (3.2) | 69.0 (10.3) | 1.7 (0.6) | 1.3 (0.6) | 83.3 (28.7) |
| L2/3 / PrL | 50 (8.2) | 29.3 (9.1) | 58 (12.6) | 19 (5.3) | 18.3 (5.1) | 96.4 (3.4) |
| L5 / PrL | 129 (12.1) | 64.3 (31.6) | 49.1 (20.3) | 97.3 (14.3) | 95 (15.7) | 97.4 (1.9) |
| L6 / PrL | 49.7 (4.0) | 27.3 (9.1) | 54.3 (13.6) | 20.7 (6.0) | 19.7 (4.5) | 96.3 (6.4) |
| L1 / IL | 1 (0) | 0.7 (0.6) | 66.7 (57.7) | 1 (1) | 1 (1) | 100 (0) |
| L2/3 / IL | 21 (9.8) | 13.3 (5.5) | 66.2 (16.3) | 3.7 (1.5) | 3.3 (1.5) | 91.7 (14.4) |
| L5 / IL | 77 (12.3) | 41.3 (26.3) | 51.1 (24.3) | 61.7 (13.6) | 60 (12.8) | 97.5 (2.5) |
| L6 / IL | 47.7 (10.2) | 31.3 (13.1) | 64.1 (15.5) | 21 (7.2) | 20.3 (7.0) | 97.1 (5.0) |

### Statistics

Principal component analysis on the densities of different GABAergic neurons across layers of PER and wS1 was performed in Matlab 2021 (Mathworks) using the function “pca”. We obtained the explained variance for each principal component, the principal component coefficients, the principal component scores, and the principal component variances. Hierarchical cluster analysis was performed in Matlab 2021 (Mathworks). After normalization with the function “zscore”, we calculated the distance between pairs of objects with the function “pdist” and the method “cityblock”. Objects were grouped according to the calculated distances with the function “linkage” and the method “average”. After calculating the inconsistency index (“inconsistent”), clusters were obtained by using the function “cluster” and the inconsistency index as cutoff. This method identified two clusters (Fig. 2A).

**Figure 2.**
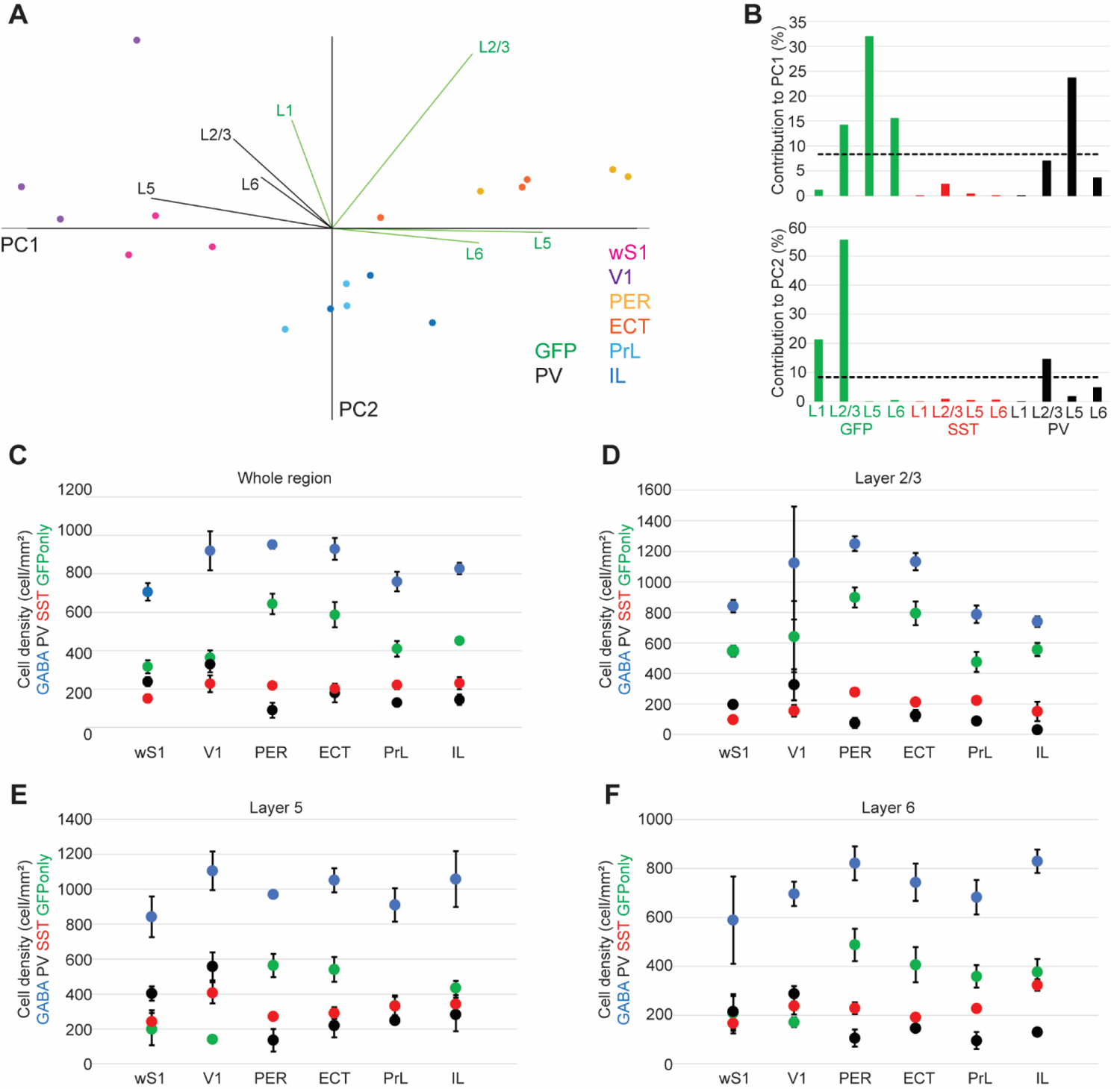
Density distribution of molecularly defined GABAergic types across six cortical areas. **A.** Data from 3 mice were plotted in principal component space using the first two principal components. Data points are color coded for cortical area and each area contains three data points representing each mouse. Vectors represent the direction and weight (length of the vector) of the variables that mostly contribute to each principal component. Vectors are color coded for cortical layer and molecular type. **B.** Bar plots showing the contribution to the variance of the population by each variable (density of molecular type in each layer) for PC1 (upper panel) and PC2 (lower panel). The dotted line shows the level at which all variables would have equal contribution. **C-F.** Plots of density of molecularly defined GABAergic neurons across cortical regions (n= 3 mice in each region). Data points are color coded for molecular type. **C.** Cell densities in whole cortical areas (i.e., combining all layers together). **D-F.** Cell densities across areas in layer 2/3 (D), layer 5 (E) and layer 6 (F).

We did not use any statistical method to pre-determine sample size. The number of mice, slices or neurons used are consistent with other studies in the field (Ma et al., 2006; Whissell et al., 2015; Kim et al., 2017; Nigro et al., 2018). No blinding method was used. When appropriate statistical comparison was performed with a one-way ANOVA with Bonferroni correction for comparing more than two groups, or with a two-sided Wilcoxon rank sum test for comparing two groups.

## Results

### Comparison of the GABAergic population of primary sensory and association areas

Previous brain-wide quantifications of cortical GABAergic neurons highlighted a prominent drop in density of PV-INs in association areas (Whissell et al., 2015; Kim et al., 2017). The interpretation of the results of these studies rests on the assumption that the mouse lines used for quantifying PV-INs reflect the intrinsic expression pattern of PV throughout the cortex (Whissell et al., 2015; Kim et al., 2017). Previous quantifications of labeled neurons in the PV-cre mouse line led to the conclusion that association areas are under a lower inhibitory control because of the low density of PV-INs (Whissel et al., 2015; Kim et al., 2017; Ding et al., 2023). However, the total GABAergic population has not yet been quantified, and other GABAergic types might take on the role of PV-INs in association areas. We have previously described that the PV-cre mouse line is not efficient in association areas (PER, ECT, LEC, PrL, IL) and captures only 50% of PV-INs (Nigro et al., 2021). To test whether the density of PV-INs is in fact lower in association areas and to quantify the total GABAergic population in these regions, we used the GAD67-GFP mouse line (n= 3) and performed immunofluorescence staining for PV, SST and GFP (Fig. 1 and 3). We found that the pattern of density distribution across cortical areas was still present when quantifying PV expression by immunofluorescence (Fig. 3A-C). PV-INs were more abundant in the dorsal sensory and motor cortices as compared to medial associative (PrL and IL) and ventro-lateral associative areas (ECT and PER) (Fig. 3A-C). We quantified the number of PV-INs, SST-INs and GFP-only neurons (neurons labeled by the GAD67-GFP mouse line but not expressing either PV or SST) across layers of six cortical areas (wS1, V1, PER, ECT, PrL, IL) and calculated their densities. We performed a PCA on the densities of these markers across the layers of these six areas to visualize in low dimensional space how these cell-types describe these six cortical regions (Fig. 2A-B).

**Figure 3.**
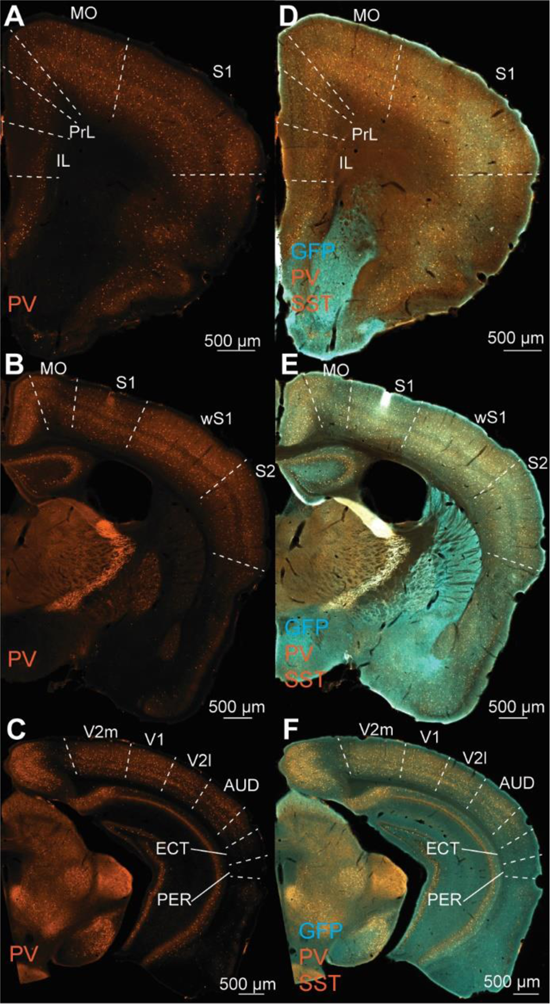
Gradients of PV expression along the rostro-caudal and dorso-ventral axes of the mouse brain. **A-C.** Representative images of a staining for PV in a GAD67-GFP mouse along the rostro-caudal axis encompassing all cortical areas included in this study. Pictures from A to C show different slices along the rostro-caudal axis. The intensity of PV staining drops in PrL/IL as compared to motor (MO) and sensory areas (S1). Proceeding caudally, the intensity of PV staining drops in lateral and ventral cortical areas (ventral to S2 in B) and at the level of PER/ECT in C. **D-F.** Same slices as in A-C but showing the signal for GFP (turquoise) and both SST and PV (orange). Note how the GFP signal becomes more evident in the areas with low PV.

The first two principal components accounted for 85.7% of the variance and the data was well segregated in the principal component space (Fig. 2A). Indeed, sensory cortices were grouped on the left side of the plot, pushed there by the higher density of PV-INs in these areas. On the other hand, association areas were spread along the horizontal axis in the right side of the PC space. The variables that best explained the variance were the density of PV-INs in layers 2/3 and 5, which was higher in sensory areas, and the density of GFP-only neurons in layers 2/3-6, which was higher in association areas (Fig. 2B).

The pattern of distribution of the density of marker-expressing neurons across the six cortical areas was in agreement with previous quantifications (Whissel et al., 2015; Kim et al., 2017) (Fig. 2C). However, the density of total GABAergic neurons was comparable across all regions (Fig. 2C). Interestingly, the density of GFP-only neurons increased in association areas, with a trend that was specular to the reduction of PV-INs in these cortices (Fig. 2C). These data suggest that a higher density of GABAergic neurons that do not express either PV or SST compensates for the reduction in PV-INs in association cortices. We found similar patterns when comparing in individual layers across cortical areas (Fig. 2 D-F).

### Electrophysiological census of the GABAergic population in wS1 and PER

Our histological examination demonstrates that although the density of PV-INs in association areas is lower than in sensory cortices, the density of GABAergic neurons is comparable. This suggests that some other GABAergic neurons might account for the increased density of GFP-only neurons. Some molecular classes of GABAergic neurons show specific biophysical properties that endow them with specific firing patterns. Indeed, PV-INs are mostly fast-spiking, whereas SST neurons are mostly adapting or low-threshold spiking (Tremblay et al., 2016). We performed an electrophysiological examination of the firing properties of GFP-expressing neurons in the GAD67-GFP mouse line (n= 18) in layer 5 of wS1 and PER. We chose wS1 because its GABAergic population has been extensively characterized, and PER because it showed the lowest density of PV-INs among the examined cortical areas.

Unexpectedly, the fast-spiking phenotype accounted for the largest population in both PER and wS1 (Fig. 4A), which is at odds with the low density of PV-INs in PER (Fig. 2). Interestingly, the proportion of fast-spiking cells was comparable in wS1 and PER (Fig. 4A). We hypothesized that some fast-spiking neurons in PER do not express detectable levels of PV. To test this hypothesis, we labeled fast-spiking neurons with a viral approach that exploits enhancer driven expression of tdTomato in fast-spiking interneurons throughout the brain (Vormstein-Schneider et al., 2020). This approach does not lean on the expression of PV in FS neurons, but the enhancer has been shown to be associated to the expression of the Nav1.1 channel (Vormstein-Schneider et al., 2020). We injected the S5E2-dTom virus in PER/ECT of C57 mice (n= 6) and characterized the electrophysiological properties of the cells labeled by the S5E2-tdTom virus (Fig. 5A-B). Most labeled neurons showed a fast-spiking phenotype (16/18), characterized by low input resistance, short action potential duration, and high maximal firing frequency with minimal adaptation (Fig. 5B). We found 2 (out of 18) neurons that showed adapting (n= 1) and LTS (n= 1) firing patterns (Fig. 5B). These firing patterns are characteristic of SST-INs as demonstrated by recordings performed in PER of SST-cre mice (n= 3 mice) (Fig. 5B).

**Figure 4.**
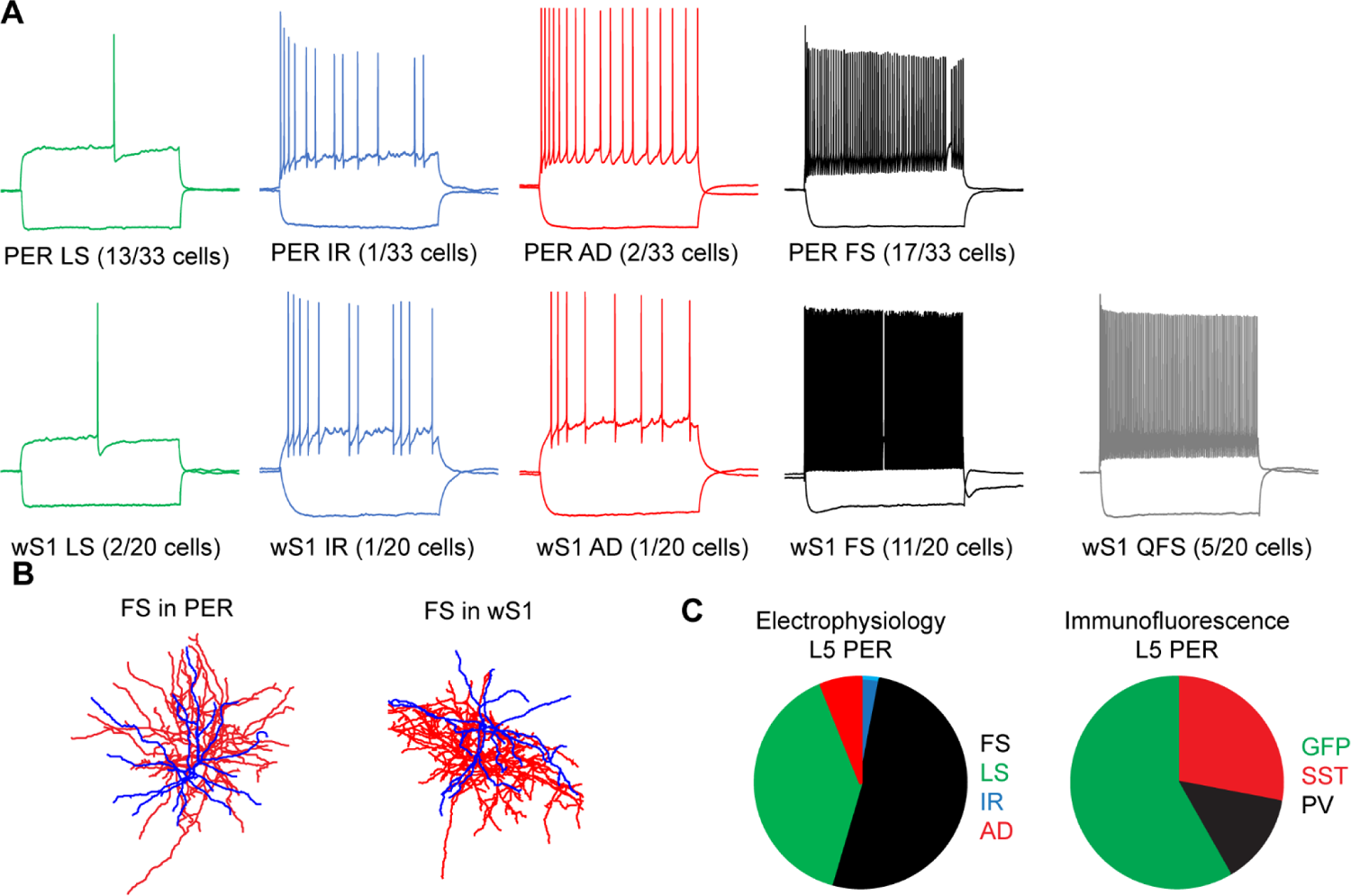
Electrophysiological census of GABAergic types in layer 5 of wS1 and PER in the GAD67-GFP mouse line. **A.** Representative voltage responses to hyper- and depolarizing current injections color coded for pattern type in PER (upper row) and wS1 (lower row). Green, late-spiking pattern characterized by a long latency to first spike near rheobase, broad AP_HW_, broad AHP and absence of sag; Blue, irregular spiking pattern characterized by early adaptation followed by spiking at irregular intervals; Red, adapting firing pattern characterized by adaptation of firing frequency followed by regular spiking; Black, fast-spiking pattern characterized by the highest maximal firing frequency, low input resistance, short AP_HW_ and fast AHP; Grey, quasi-fast spiking pattern (only in wS1), similar to the fast spiking but with higher IR, lower firing rates and stronger adaptation (see Ma et al., 2006; Nigro et al., 2018). **B.** Representative recovered morphologies of fast-spiking neurons in PER (left) and wS1 (right). Dendrites are colored in blue and axons in red. **C.** Pie charts comparing the percentages of different firing patterns (left) and molecular types (right) in layer 5 of PER. Color codes follow the one used for firing pattern in A and assumes the equivalence of the fast-spiking pattern with PV-INs.

**Figure 5.**
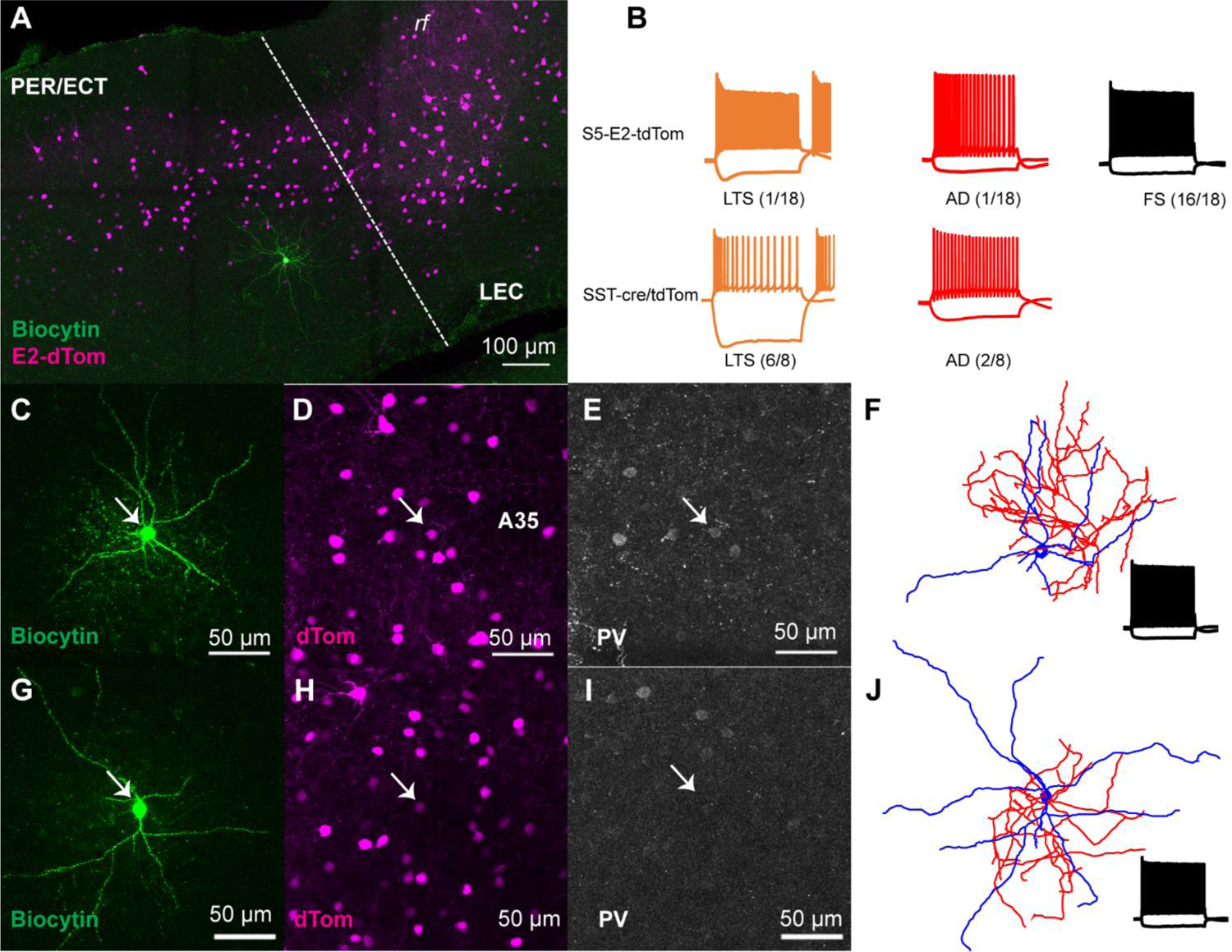
Electrophysiological characterization of neurons labeled by the S5E2-dTom virus in layer 5 of PER of C57 mice. **A.** Representative image of a brain slice containing PER from a C57 mouse injected with the S5E2-dTom virus. Neurons labeled by the virus are shown in magenta and the recorded neuron in green. LEC, lateral entorhinal cortex; rf, rhinal fissure. **B.** Representative voltage responses to hyper- and depolarizing current injections of neurons recorded from C57 mice injected with the S5E2-dTom virus (upper row) and in the PER of SST-cre/tdTom mice. Voltage responses are color coded for firing patterns: orange, low-threshold spiking characterized by high input resistance, low threshold for AP generation and rebound bursting (Ma et al., 2006; Nigro et al., 2018); red, adapting firing pattern; black, fast-spiking pattern. **C-E.** Example neurons showing the expression of PV in fast-spiking neurons labeled by the S5E2-dTom virus in PER. **F.** Recovered morphology and firing pattern of the cell in C. Dendrites are colored in blue and axons in red. **G-I.** Lack of PV expression in an example fast-spiking neuron labeled by the S5E2-dTom virus in PER. **J.** Recovered morphology and firing pattern of the neuron in G.

To directly test the hypothesis that some fast-spiking neurons in PER do not express PV, we performed immunofluorescence for PV on 15 of the recorded fast-spiking neurons and found that 6 out of 15 did not express PV (E2-PV-) (Fig. 5C-I). A comparison of the electrophysiological properties of PV-IN and E2-PV-revealed that the two populations have similar biophysical properties despite the difference in PV expression (Table 2). Interestingly, when comparing fast-spiking neurons in PER and wS1, we found several electrophysiological differences, with fast-spiking neurons in wS1 having faster HW and AHPs, which enables them to reach a higher maximal firing frequency (Fig. 6B-G). We used hierarchical clustering on the electrophysiological parameters and further confirmed the difference between fast-spiking neurons in the two cortical areas (Fig. 6A). In PER PV-INs and E2-PV-were clustered together, further confirming that their electrophysiological properties are indistinguishable (Fig. 6A).

**Figure 6.**
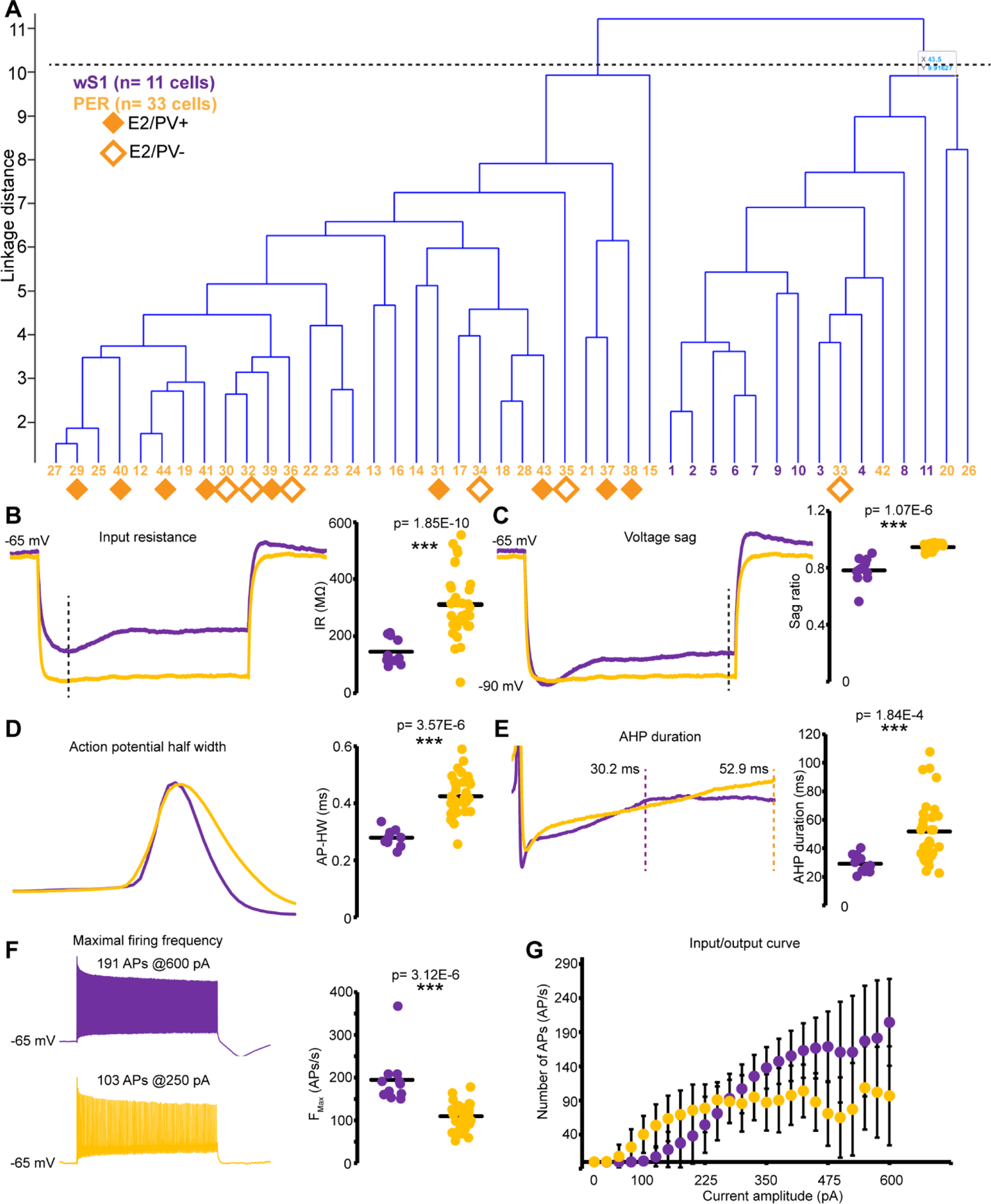
Fast-spiking neurons show region-specific electrophysiological properties independently of PV expression. **A.** Dendrogram obtained by hierarchical cluster analysis of electrophysiological properties of neurons in wS1 and PER. The cluster analysis revealed two clusters that largely overlap with the area of where neurons were recorded rather than with the expression of PV. **B-G.** Comparison of the significantly different electrophysiological properties between wS1 (purple) and PER (yellow) neurons. Each panel shows representative voltage responses showing the indicated electrophysiological feature (left panel) and the plot of the distribution for all neurons in the two areas (right panel). See Methods for details on the measurements of each electrophysiological property. p values for a Wilcoxon rank sum test are shown for each plot.

**Table 2.**
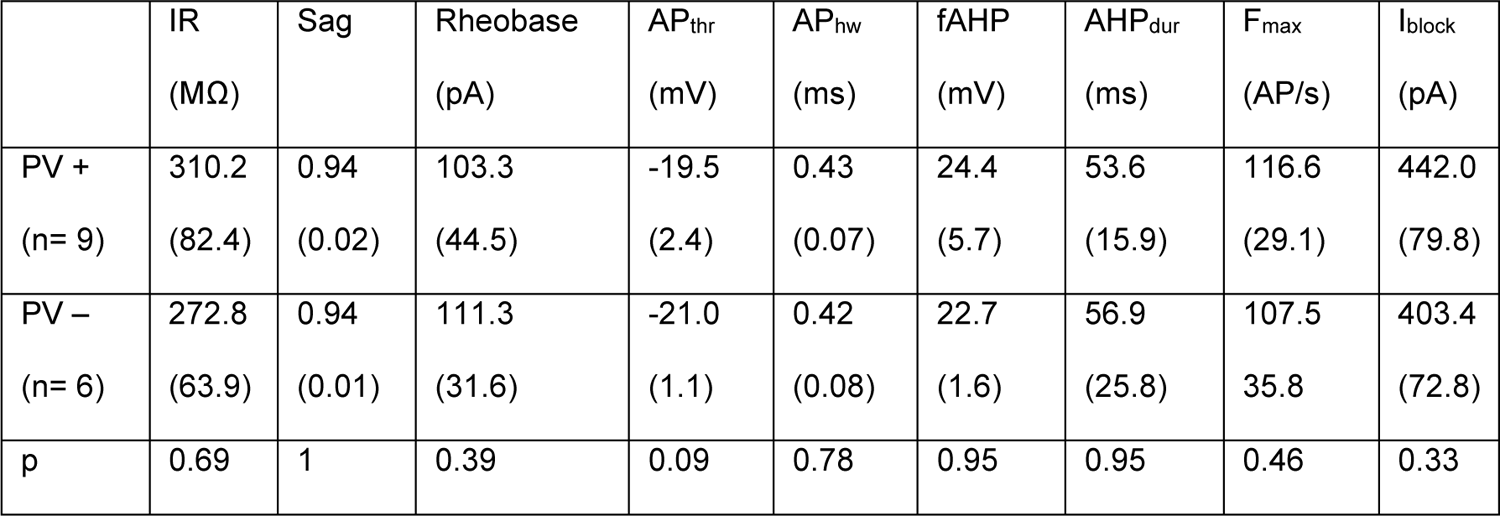
Electrophysiological properties of fast-spiking cell expressing PV and not expressing PV in layer 5 of PER. Cells were labeled with the S5E2-dTom virus injected in C57 mice. Values report the mean and standard deviation (in parentheses). The last row reports the p value of a Wilcoxon rank sum test.

|  | IR<br>(MΩ) | Sag | Rheobase<br>(pA) | AP<sub>thr</sub><br>(mV) | AP<sub>hw</sub><br>(ms) | fAHP<br>(mV) | AHP<sub>dur</sub><br>(ms) | F<sub>max</sub><br>(AP/s) | I<sub>block</sub><br>(pA) |
| --- | --- | --- | --- | --- | --- | --- | --- | --- | --- |
| PV +<br>(n= 9) | 310.2<br>(82.4) | 0.94<br>(0.02) | 103.3<br>(44.5) | -19.5<br>(2.4) | 0.43<br>(0.07) | 24.4<br>(5.7) | 53.6<br>(15.9) | 116.6<br>(29.1) | 442.0<br>(79.8) |
| PV –<br>(n= 6) | 272.8<br>(63.9) | 0.94<br>(0.01) | 111.3<br>(31.6) | -21.0<br>(1.1) | 0.42<br>(0.08) | 22.7<br>(1.6) | 56.9<br>(25.8) | 107.5<br>(35.8) | 403.4<br>(72.8) |
| p | 0.69 | 1 | 0.39 | 0.09 | 0.78 | 0.95 | 0.95 | 0.46 | 0.33 |

Our electrophysiological characterization of GABAergic neurons demonstrates that PER contains a large population of fast-spiking neurons that do not express PV and are not labeled by the PV-cre mouse line. These neurons though can be labeled by an enhancer-driven viral strategy that is independent from the expression of PV. Does this population of fast-spiking neurons not expressing PV account for the increased proportion of GABAergic neurons not expressing PV or SST in association cortices?

### Association cortices contain fast-spiking interneurons that do not express PV

The experiments described above demonstrate that PER contains a large number of fast-spiking GABAergic neurons that do not express PV, and that has been so far unacknowledged by previous estimations of GABAergic types across cortical areas (Whissell et al., 2015; Kim et al., 2017). To test whether this population is also present in other association areas, we developed an intersectional viral approach to specifically label fast-spiking GABAergic neurons while excluding SST-INs (Fig. 7). As demonstrated by our electrophysiological characterization of the E2-dTom labeled neurons, we found that SST-INs account for about 20% of the labeled neurons in PER/ECT of GAD67-GFP mice (n= 2) (Fig. 7A-C). To exclude the SST-INs, we inserted lox sites for the cre-dependent excision of the tdTomato sequence and additional lox sites for the cre-dependent inversion of an inverted sequence coding for GFP (Fig. 7D). When injected in PER/ECT of an SST-cre mouse, this strategy labeled E2/SST-INs with GFP and E2/SST-fast-spiking neurons with tdTomato (Fig. 7E-G). We found virtually no overlap between E2-dTom and E2-GFP neurons (Fig. 7F) and virtually all E2-GFP neurons expressed SST (Fig. 7G). To test whether fast-spiking neurons not expressing PV are present in other association areas, we injected the intersectional virus in PER/ECT and PrL/IL cortex of SST-cre mice (n= 2) (Fig. 8A-B). In PER/ECT, only about 40% of putative fast-spiking neurons express PV, while about 65% of putative fast-spiking neurons in PrL/IL express PV, suggesting that a large population of fast-spiking neurons in association cortices do not express PV (Fig. 8C). We also quantified the density of tdTomato expressing neurons in PER/ECT and in PrL/IL areas and observed that it was comparable to the density of PV-INs in wS1 and V1 (Fig. 8D), confirming our hypothesis that the population of fast-spiking neurons not expressing PV accounts for the reduced density of PV-INs previously reported in these areas (Whissell et al., 2015; Kim et al., 2017). It was hypothesized that the reduced density of PV-INs in association areas might be followed by a reduction of their output and a higher excitability of the network, leading to enhanced recurrency (Kim et al., 2017; Ding et al., 2023). We tested this hypothesis by injecting an intersectional virus carrying a loxP flanked Channelrhodopsin (ChR2) sequence in PER of SST-cre mice (n= 1 mouse). This virus allows the expression of ChR2 only in putative fast-spiking neurons while excluding SST-INs (Fig. 9). We obtained acute slices of PER/ECT and performed voltage clamp experiments to measure the light evoked IPSC from layer 5 excitatory neurons in this region (Fig. 9A). When comparing these results with those obtained from wS1 in PV-cre/ChR2 mice, we found that the IPSCs amplitude was not significantly different from that in PER (wS1: 12.2 ± 4.8 nA, n= 6 cells; PER-E2: 7.9 ± 2.8 nA, n= 8 cells; p= 0.09, Bonferroni correction) (Fig. 9). These results demonstrate that the net output of fast-spiking neurons is not significantly different in the two areas, further confirming that the fast-spiking population is comparable across cortical regions independently from PV expression.

**Figure 7.**
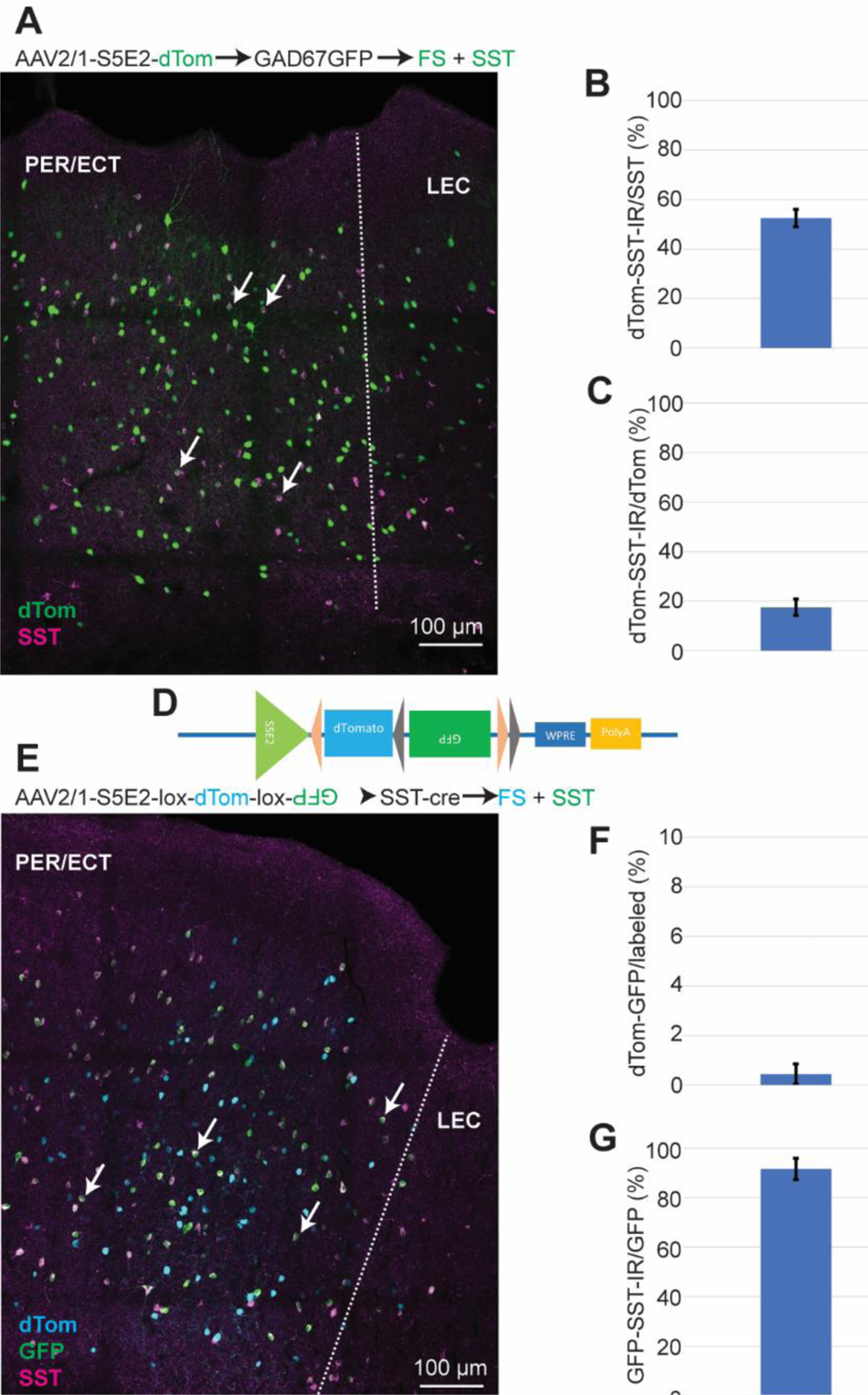
Intersectional viral strategy to label fast-spiking neurons while excluding SST-INs. **A.** Representative slice of PER/ECT of a GAD67-GFP mouse injected with the S5E2-dTom virus. Immunofluorescence for SST (magenta) and dTom (green) shows that some neurons labeled by the virus express SST (arrows). **B.** Plot showing the percent of SST-INs labeled by the S5E2-dTom virus (n= 2 mice). **C.** Plot showing the percent of neurons expressing dTom (labeled by the virus) that also express SST (n= 2 mice). **D.** Schematics of the sequence of the intersectional virus. Pinks triangles represent lox 2272 sites and gray triangles represent loxP sites. **E.** Representative slice of PER/ECT of a SST-cre mouse injected with the intersectional virus. Immunofluorescence for dTom (turquoise), GFP (green) and SST (magenta) shows no overlap between dTom and GFP expression and SST expression only in GFP labeled neurons (arrows). LEC, lateral entorhinal cortex. **F.** Plot showing the percent of neurons expressing dTom and GFP in the population of labeled neurons (dTom + GFP) (n= 2 mice). **G.** Plot showing that virtually all GFP neurons express SST (n= 2 mice).

**Figure 8.**
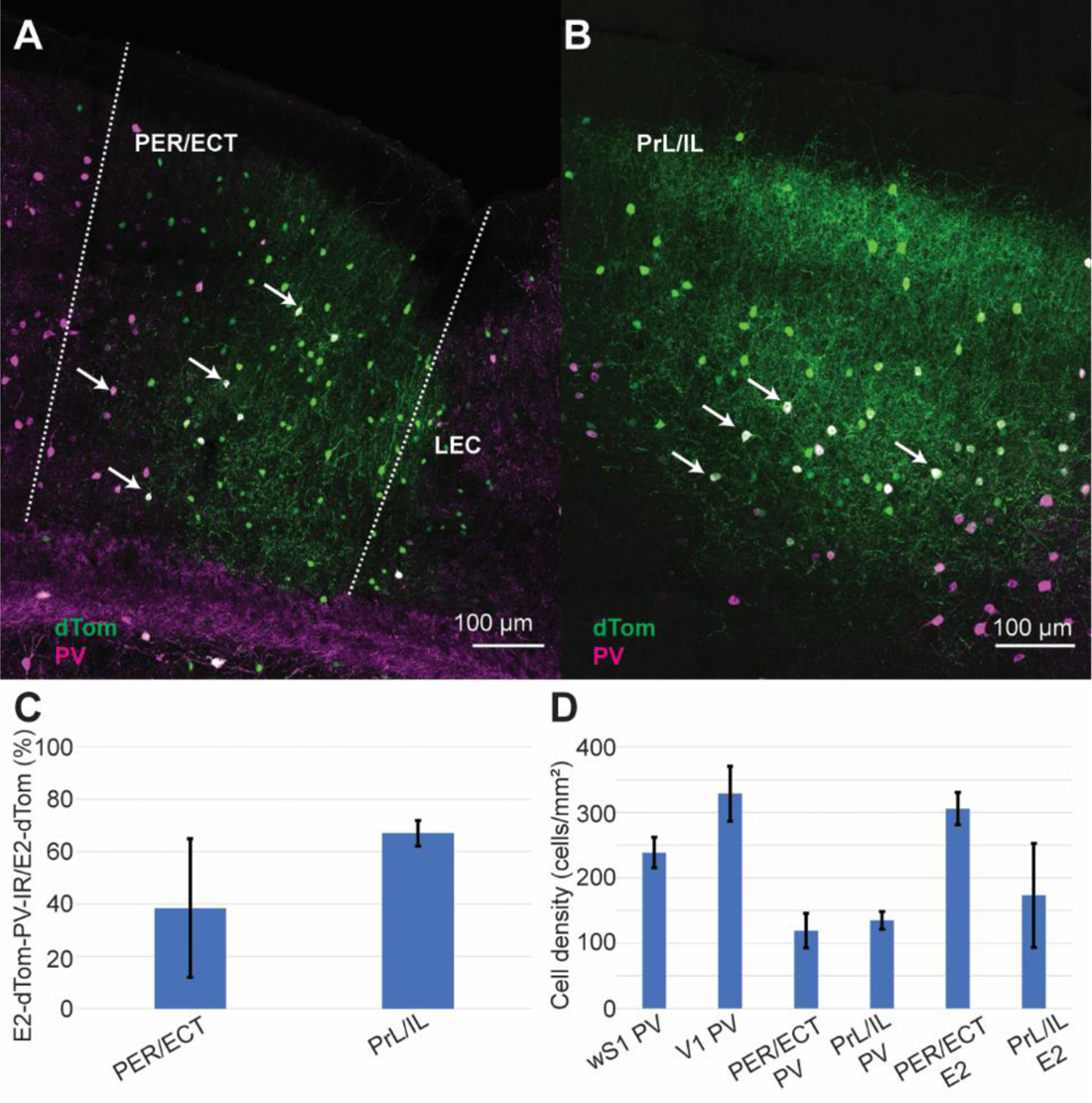
Fast-spiking neurons that do not express PV are present in multiple association cortices. **A-B.** Representative slices of PER/ECT (A) and PrL/IL (B) of SST-cre mice injected with the intersectional virus. Immunofluorescence for PV (magenta) revealed that some dTom (green) neurons do not express PV. Arrows indicate cells labeled by the virus and expressing PV. **C.** Plot showing the percent of dTom expressing neurons that also express PV in PER/ECT and PrL/IL (n= 2 mice). **D.** Bar plot comparing the density of putative fast-spiking neurons across cortical regions inferred from PV expression (first four columns, as in Fig. 1C) and from the labeling by the intersectional virus (last two columns).

**Figure 9.**
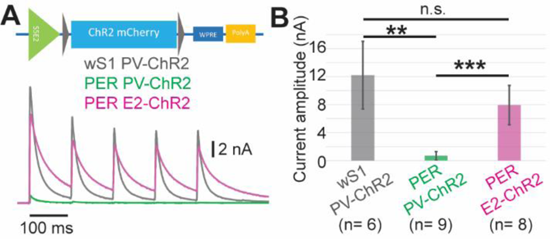
The magnitude of the synaptic output of fast-spiking neurons is equivalent across cortical regions. **A.** Top, schematics of the sequence of the intersectional optogenetic virus. Bottom, representative voltage clamp recordings of outward currents evoked by optogenetic activation of neurons labeled in the PV-ChR2 mouse line or by the intersectional method. Note how the intersectional strategy allows to recover a much larger current in PER as compared to the PV-ChR2 mouse line. **B.** Statistical comparison of the amplitude of synaptic currents evoked in wS1 and PER showing not statistically significant difference between wS1 and PER when the intersectional virus is used to target all fast-spiking neurons. Statistical comparison used a one-way ANOVA with Bonferroni correction: n.s., p= 0.09; two asterisks, p= 0.002; three asterisks, p= 0.0001. Number of neurons in parentheses.

## Discussion

The present study challenged recent findings showing that the density and functional output of fast-spiking neurons in association cortices is lower than that of sensory cortices (Whissell et al., 2015; Kim et al., 2017). We demonstrate that association cortices contain a comparable density of GABAergic neurons to sensory areas despite the low density of PV-INs. We provide multiple lines of evidence demonstrating that the fast-spiking population of association cortices is comparable to that of wS1, and a considerable number of these neurons do not express PV. We developed an intersectional viral strategy to label fast-spiking neurons, including those not expressing PV in association areas, and used this tool to demonstrate that the net output of fast-spiking neurons on the excitatory population is comparable across cortical areas.

The low expression of PV in association cortices has been used by neuroanatomists to delineate these areas for several decades in different species (Pitkänen and Amaral, 1993; Burwell et al., 1995; Uva et al., 2004; Van De Werd et al., 2010; Beaudin et al., 2013). However, this observation has only recently been examined in the context of GABAergic cell-types and inhibitory circuits using specific mouse lines targeting PV-INs (Whissel et al., 2015; Kim et al., 2017). We have recently described that the labelling in the PV-Cre mouse line is very inefficient and quantifications of the number of PV-INs would be dramatically underestimated (Nigro et al., 2021). The present study corroborates the low density of PV-INs in PER/ECT by quantifying PV-INs neurons and extends this finding to PrL/IL areas. The fast-spiking population not expressing PV has never been reported previously, likely because of the absence of a labeling tool, such as a molecular marker or mouse line. However, these neurons can be labeled by a viral strategy that labels all fast-spiking neurons and that is independent of PV expression. This further suggests that the two fast-spiking populations share other features related to gene expression.

Our electrophysiological characterization demonstrates that fast-spiking interneurons show different biophysical properties in wS1, and PER. This might reflect specializations of biophysical properties related to computations performed in these two brain regions. In wS1 (and other primary sensory areas) PV-INs are involved in fast feedforward inhibition to constrain the integration window of excitatory postsynaptic potentials (Tremblay et al., 2016). Previous studies have shown that a large percent of excitatory neurons in PER/ECT show a long latency for spike generation that might contribute to gating of inputs and to temporal integration (Storm, 1988; Faulkner and Brown, 1999; Beggs et al., 2000; Kajiwara and Tominaga, 2021). These features of the excitatory population might not require the fast inhibitory dynamics typical of sensory region and that are allowed by the membrane properties that characterize fast-spiking neurons in sensory cortices.

Variations of the morpho-electrical properties of neuron types and their circuits have been described within cortical areas (Fletcher and Williams, 2019; Large et al., 2018). Other studies have highlighted the distribution of specific cell types across cortical areas, pinpointing at regional specializations (Scala et al., 2019). These studies and our results suggest the importance of investigating the biogeography of cortical neurons to understand the circuits underlying region-specific computations.

The expression of PV in the cortex increases throughout the first three postnatal weeks (Goldberg et al., 2011), and it has been shown to be activity dependent (Donato et al., 2013). All our experiments were performed on adult mice (> 2 month), a time when the pattern of PV expression has already been established. Upregulation of PV in neurons with low levels of PV expression has been demonstrated in the hippocampus of mice upon training on a task (Donato et al., 2013). However, it has never been assessed whether this occurs in GABAergic neurons that do not express PV. Future studies should address how network activity interacts with PV expression in the two populations of fast-spiking neurons in association areas. Interestingly, rat models of temporal lobe epilepsy show a reduced number of PV-INs (Benini et al., 2011). The loss of PV expression in epileptic tissue might be due to a loss of expression of PV rather than to cell death of PV-INs (Filice et al., 2016; Medici et al., 2016). The lack of PV expression in a large population of fast-spiking neurons in association areas suggests that these areas might be more prone to seizures (Biagini et al., 2013). The intersectional strategy described here could provide a tool to manipulate the activity of these neurons in epilepsy models to unveil their contribution to seizure generation.

Modeling studies showed that the density of PV-INs across cortical areas is tightly correlated with a connectivity based hierarchical organization of the cortex (Kim et al., 2017; Ding et al., 2023). These studies assumed that the low density of fast-spiking neurons in association areas and the circuit motifs of the cortex promote disinhibition of the excitatory population and increased recurrency in these cortices (Kim et al., 2017; Ding et al., 2023). However, our results demonstrate that not only is the density of fast-spiking neurons comparable across cortical areas, but so is their synaptic output onto excitatory neurons, suggesting that other mechanisms might underly recurrency in association cortices. These modeling studies also suggest that inhibition from SST-INs onto fast-spiking neurons is fundamental in establishing recurrent activity in association cortices. Future studies might test whether fast-spiking neurons in association cortices receive stronger inputs from SST-INs, which might promote recurrency as suggested by previous modeling studies (Kim et al., 2017; Ding et al., 2023).

## Acknowledgments

We thank Dr. Grethe Olsen for excellent technical assistance and all members of the Nigro group at the Kavli Institute for Systems Neuroscience for constructive criticism. We also thank Dr. Bernardo Rudy, Dr. Robert Machold and Dr. Hector Zurita for their comments on an earlier version of this manuscript. fThis research was funded by the Norwegian Research Council through the Center of Excellence scheme – Center for Neural Computation (Grant No. 223262 to MPW) and Center for Algorithms in the Cortex (Grant No. 321969 to MJN), the European Union’s Horizon 2020 Research and Innovation Programme under the Marie Skłodowska-Curie (Grant No. 885955 to MJN) and the Kavli Foundation.

## Author contribution

MJN designed research; EJC, KK, LC, MJN performed research; EJC, KK, MJN analyzed data; RRN contributed unpublished reagents; MPW and MJN provided funding; MJN supervised research; MJN wrote the paper with input from all authors.

